# Trophic specialization on unique resources in one of the most celebrated examples of sympatric speciation, Barombi Mbo crater lake cichlids

**DOI:** 10.1101/2021.08.04.455125

**Authors:** Jacquelyn R. Galvez, Keara McLean, Cyrille Dening Touokong, Legrand Nono Gonwouo, Christopher H. Martin

## Abstract

Divergent ecological selection often results in trophic niche partitioning and is one of the central processes underlying sympatric speciation. However, there are still few studies of niche partitioning in putative examples of sympatric speciation in the wild. Here we conducted the first quantitative study of dietary niche partitioning in one of the most celebrated examples of sympatric speciation, Barombi Mbo cichlids, using stomach contents and stable isotope analyses. We found little evidence for trophic niche partitioning among any Barombi Mbo cichlids, even among the nine species coexisting in sympatry in the littoral zone. Stable isotope analyses supported these conclusions of minimal dietary overlap. However, we did find extraordinary dietary specialization in some species, including spongivory and feeding on terrestrial ants, both unique feeding strategies among freshwater fishes. Stomach contents of the spongivore (*Pungu maclareni*) were 20% freshwater sponge, notable considering that only 0.04% of all fishes consume sponges. Overall, we conclude that while there is less trophic niche partitioning than expected among Barombi Mbo cichlids, there is evidence for dietary specialization on rare resources in support of Liem’s paradox.

## INTRODUCTION

All adaptive radiations display some level of niche partitioning, a phenomenon in which groups of organisms in a shared environment shift their resource use to reduce niche overlap. While phylogenetic niche conservatism results in ecological similarities among closely related species (McNyset 2009, Losos 2008), adaptive radiations represent the opposite extreme in which rapidly diversifying species occupy a diverse suite of ecological niches (Schulter 2000, Martin & Richards 2019, Stroud & Losos 2020). Trophic niche partitioning in particular can allow for the coexistence of similar species by reducing interspecific competition for limited food resources (Ross 1986, Winemiller & Pianka 1990, Correa & Winemiller 2014, Varghese *et al.* 2014). When such niche shifts coincide with traits under disruptive selection and assortative mating, dietary niche partitioning can contribute to ecological speciation in sympatry (Dieckmann and Doebeli 1999, Gavrilets 2004, Gavrilets & Losos 2009). Examples of this phenomenon include experimental evolution studies (Blount *et al.* 2008), divergence of insect host races (Nosil 2009), and ecological speciation in classic adaptive radiations (Grant and Grant 2002; Kocher et al. 2004; Lamichhaney et al. 2015; Gillespie *et al.* 2020). The fine-scale study of dietary niche partitioning and specialization during adaptive radiation can offer further insight into the prevalence and mechanisms of the processes driving ecological speciation.

African cichlids are widely regarded as a model system for studying adaptive radiation. The species flocks of Lakes Malawi, Victoria, and Tanganyika in particular contain dietary specialists and closely related species that exhibit varying levels of trophic niche partitioning (Kocher 2004; Martin and Genner 2009; Wagner *et al.* 2009). Despite this, dietary specialization is rarely invoked as the driver of speciation in African cichlids. Habitat partitioning (Albertson 2008; Conith *et al.* 2020) and sexual selection (Seehausen 2000; Poelstra *et al.* 2018) are hypothesized to play a much greater role in the observed ecological diversity of these lineages. In fact, there is considerable dietary overlap among many sympatric rock-dwelling Malawi cichlids (Ribbink et al. 1983; Reinthal 1990; Genner *et al.* 1999*a*; Genner *et al.* 1999b; Martin & Genner 2009), suggesting that many closely related cichlid species can coexist without strong ecological segregation. It has long been recognized in Lake Malawi that closely related species often show minimal or undetectable ecological differentiation, despite substantial differences in trophic morphology (Liem 1980). This is known as Liem’s paradox: trophic specialists act as “jacks-of-all-trades” able to consume both their narrow food source as well as a more generalist diet (Liem 1980). A rare exception is observed in the *Alcolapia* species flock of the alkaline Lake Natron (Seegers & Tichy 1999). These three herbivorous species exhibit fine-scale trophic and ecomorphological differences despite limited genomic differentiation, suggesting the importance of trophic niche partitioning in the diversification of this clade (Ford *et al.* 2016).

Another African cichlid radiation displaying evidence of trophic niche partitioning is in crater lake Barombi Mbo, Cameroon. This species flock of 11 endemic cichlids is celebrated as a putative example of sympatric speciation in nature and its discovery led to a revival of empirical and theoretical interest in this process (Turelli *et al.* 2001; Schliewen *et al.* 1994; Schliewen and Klee 2004; Coyne & Orr 2004; Bolnick & Fitzpatrick 2007; Richards *et al.* 2019). However, recent work on this system has revealed a complex history of gene flow between Barombi Mbo cichlids and riverine outgroups (Martin *et al.* 2015), with introgression from multiple colonizations potentially contributing to the speciation process (Richards *et al.* 2018).

More directly relevant to niche partitioning, there is also evidence for weak disruptive selection within Barombi Mbo cichlids (Martin 2012). Strong disruptive selection is necessary in all theoretical models of sympatric speciation to drive the evolution of reproductive isolation between ecotypes (Dieckmann & Doebeli 1999; Gavrilets 2004; Otto et al. 2008). Compared to the predicted strength of disruptive selection necessary for sympatric speciation (Dieckmann and Doebeli 1999; Bolnick 2011) and empirical estimates of disruptive selection in nature (Kingsolver *et al.* 2001), cichlids of the Barombi Mbo genus *Stomatepia* displayed relatively weak levels of disruptive selection for all trophic morphology traits measured (Martin 2012). While trophic divergence in *Stomatepia* has been previously reported for stable isotope data (Martin 2012), divergence in overall dietary profiles—i.e. stomach contents—has not been assessed since Trewavas *et al.* (1972). Unlike the cichlids of Cameroonian Lake Ejagham—another endemic cichlid lake radiation recognized as an example of sympatric speciation—in which olfactory preferences and sexual selection are hypothesized to drive divergence (Martin 2013; Poelstra *et al.* 2018), striking differences among sympatric species in trophic morphology and no sexual dimorphism in ten out of the eleven species in Barombi Mbo suggest that diet could be the primary driver of ecological speciation. Previous qualitative descriptions of diet indicated differences in dietary profiles among some species and identified species that fed heavily on plants and freshwater sponges (Trewavas *et al.* 1972).

In this study, we investigated patterns of trophic niche partitioning and specialization among Barombi Mbo cichlids, namely through stomach content and stable isotope analyses. We used stomach content analyses to quantify differences in dietary item proportions, niche width and overlap, and overall dietary composition. We also used stable isotope analyses to investigate relative trophic levels and carbon source differences among species over a longer timeframe than the “snapshot” provided by stomach content data. Investigating dietary and trophic differences and specialization among Barombi Mbo cichlids is the first step in examining whether diet is the primary driver of sympatric speciation in this system.

## METHODS

### Study site and sample collection

Barombi Mbo is a 1 Mya volcanic crater lake (Cornen *et al.* 1992) in southwestern Cameroon. It is roughly circular in shape with a diameter of 2.5 km and a maximum depth of 110 m, but the oxic zone only reaches to 30 m (Trewavas *et al.* 1972; Cornen *et al.* 1992; Musilova *et al.* 2019). We collected samples in December 2009 through January 2010, and in July through December, 2016 from several localities in the lake using a 6 x 2 m seine net with 0.5 cm^2^ mesh. *Sarotherodon linnellii* and *Konia dikume* were caught by artisanal fishers using gill nets. We collected all 11 endemic Barombi Mbo cichlid species. We euthanized captured fish with an overdose of MS-222 and immediately took a 5 mg muscle tissue sample from the caudal peduncle for stable isotope analysis. Muscle samples were desiccated individually with magnesium perchlorate in airtight vials following Martin (2012; 2013). Specimens were then individually labeled and fixed in 95-100% ethanol. Field procedures followed approved protocols by the Institutional Animal Care and Use Committees of the University of California, Davis and the University of North Carolina at Chapel Hill.

### Stomach content analyses

In total, we selected 241 individuals for stomach content analysis, including at least 8 individuals from each species. Nine out of the lake’s 11 endemic cichlid species were analyzed in this study, all except *Sarotherodon caroli* and *Sarotherodon lohbergeri*, which are morphologically and ecologically similar to *Sarotherodon steinbachi*. We removed the entire stomach and intestine from each individual. We then placed stomach contents or a subset of the intestine on a Sedgwick-Rafter cell containing 1 x 1 mm squares for visualization and quantification under a stereomicroscope. Dietary proportions were based upon a visual volume estimation method (Hyslop 1980; Manko 2016). We compressed stomach contents to a uniform thickness (approx. 0.5 mm) and estimated the surface area of each prey item by counting the number of 1 mm^2^ squares covered by the item (Hyslop 1980; Gelwich & McIntyre 2017). Smaller items were assigned fractions of a square to the nearest 0.1 mm^2^. This number was then divided by the total number of squares covered by all diet items for that individual to calculate individual dietary proportions for each item. Proportions were rounded to the nearest 0.001 and are reported as percentages.

We identified all diet items based on partially digested remnants, including exoskeletal remains, plant matter, and sponge spicules; unidentified organic matter was classified as detritus and inorganic matter, such as particles of sand, was classified as silt. All prey items were grouped into taxa, usually to the level of class or family. Diet categories were comparable to previously identified prey items of Barombi Mbo cichlids described in Trewavas *et al.* (1972). We used 13 diet categories in total: ants, *Corvospongilla spp.* sponge, Dipteran larvae, Ephemeropteran larvae, Trichopteran larvae, fish, gastropod shell, nematode, plant tissue, shrimp, detritus, silt, and unidentified. Ants were identified by distinct head capsules of species within Formicidae, which likely originated from terrestrial debris that fell into the lake. The sponge category consisted of two members of the genus *Corvospongilla*: *C. thysi*, endemic to Barombi Mbo, and closely related *C. bohmii* (Trewavas *et al.* 1972). Both species are found in the lake’s shallow waters (up to 3-4 m depth), with *C. thysi* typically covering the outer surfaces of rocks and *C. bohmii* found in crevices (Trewavas *et al.* 1972). Dipteran larvae included larval forms of the midge families Chaoboridae and Chironomidae. Ephemeropteran larvae included larval forms of various mayfly families Baetidae and Caenidae. This category also included larvae of the common burrowing mayfly species *Povilla adusta*, previously identified by Trewavas *et al.* (1972) to be present on both stones and fallen logs in Barombi Mbo. Trichopteran larvae consisted of caddisflies in their larval form, likely from the genus *Triaenodes*, which has many species endemic to West Africa (Andersen & Holzenthal 2002). The fish category was assigned to portions of fish fins and tissue, as well as to whole fry found in individuals’ stomachs (not identifiable to the species level at this size). The gastropod shell category consisted of shell remains from various snails, including freshwater limpets from the genus *Ferrissia* (Trewavas *et al.* 1972). The nematode category contained all roundworms, likely including both terrestrial and aquatic species. The plant tissue category was assigned to all plant material found in individuals’ stomachs. This included aquatic species such as *Najas pectinate* and *Potamogeton octandrus* previously documented in Barombi Mbo (Trewavas *et al.* 1972) and any terrestrial plant leaves. The shrimp category consisted of *Caridina spp.* and *Macrobranchium spp.*, freshwater shrimp genera found in Barombi Mbo and throughout Cameroon (Trewavas *et al.* 1972). Detritus was used as a catch-all term to describe organic matter that was digested beyond the point of identification. Silt was used as a catch-all to describe inorganic materials, including rocks and sand. Animal remains that could not be clearly identified (e.g. egg-like structures) were grouped into the unidentified category.

We estimated dietary niche breadth of each species by calculating Levins’ standardized index (Levins 1968) and Pianka’s measure of dietary niche overlap (Pianka 1973) using the *spaa* package (Zhang 2016) in R (version 4.0.2). Individuals with empty stomachs were excluded from all calculations (*n* = 38).

### Stable isotope analyses

To assess relative trophic positions of Barombi Mbo cichlids, we performed stable isotope analyses for all 11 species (including *S. caroli* and *S. lohbergi*). In limnetic systems, δ13C isotope ratios offer insight into the ultimate carbon source of prey consumed (Post 2002). Higher δ13C values indicate a more littoral carbon source, while lower values indicate a more pelagic source (Post 2002). δ15N ratios indicate the relative trophic position of individual consumers (Post 2002). In total, we selected 180 individuals for stable isotope analysis, including at least 6 individuals from each species. Field samples desiccated with magnesium perchlorate in individual vials were subsequently dehydrated at 60° C for at least 24 hours, then 1 mg samples were weighed to the nearest 0.0001 g, packaged into tinfoil capsules, and sent to the UC Davis Stable Isotope Facility. ^13^C and ^15^N isotopes were measured on a PDZ Europa ANCA-GSL elemental analyzer interfaced to a PDZ Europa 20–20 isotope ratio mass spectrometer (Sercon Ltd., Cheshire, U.K.).

### Statistical analyses

Individuals with empty stomachs (*n* = 38) were excluded from all statistical analyses of stomach contents, leaving a final sample size of *n* = 203. The sample size for each species is reported in Table 1. We performed all statistical analyses in R version 4.0.2 (R Core Team 2020).

**Table 1:** Mean proportion of each dietary component and sample sizes by species. BA is Levins’ standardized index of niche breadth (Levins 1968).

| Dietary component | *Konia dikume* | *Konia eisentrauti* | *Myaka myaka* | *Sarotherodon linnellii* | *Sarotherodon steinbachi* | *Pungu maclareni* | *Stomatepia mongo* | *Stomatepia mariae* | *Stomatepia pindu* |
| --- | --- | --- | --- | --- | --- | --- | --- | --- | --- |
| Ant | 0.000 | 0.000 | 0.000 | 0.000 | 0.000 | 0.000 | 0.000 | 0.098 | 0.000 |
| *Corvospongilla* Sponge | 0.000 | 0.000 | 0.000 | 0.000 | 0.000 | 0.212 | 0.000 | 0.000 | 0.000 |
| Detritus | 0.889 | 0.628 | 1.000 | 0.865 | 0.608 | 0.609 | 0.143 | 0.526 | 0.651 |
| Dipteran Larvae | 0.111 | 0.027 | 0.000 | 0.000 | 0.000 | 0.003 | 0.000 | 0.023 | 0.000 |
| Ephemeropteran Larvae | 0.000 | 0.101 | 0.000 | 0.025 | 0.009 | 0.088 | 0.000 | 0.120 | 0.000 |
| Fish | 0.000 | 0.000 | 0.000 | 0.060 | 0.000 | 0.000 | 0.000 | 0.074 | 0.000 |
| Gastropod Shell | 0.000 | 0.000 | 0.000 | 0.000 | 0.000 | 0.003 | 0.000 | 0.000 | 0.000 |
| Nematode | 0.000 | 0.000 | 0.000 | 0.000 | 0.000 | 0.000 | 0.000 | 0.015 | 0.000 |
| Plant Tissue | 0.000 | 0.228 | 0.000 | 0.045 | 0.000 | 0.016 | 0.019 | 0.006 | 0.000 |
| Shrimp | 0.000 | 0.000 | 0.000 | 0.000 | 0.000 | 0.008 | 0.629 | 0.001 | 0.091 |
| Silt | 0.000 | 0.012 | 0.000 | 0.000 | 0.000 | 0.002 | 0.000 | 0.000 | 0.000 |
| Trichopteran Larvae | 0.000 | 0.000 | 0.000 | 0.000 | 0.000 | 0.000 | 0.000 | 0.000 | 0.141 |
| Unidentified | 0.000 | 0.005 | 0.000 | 0.006 | 0.383 | 0.059 | 0.209 | 0.137 | 0.117 |
| n | 3 | 37 | 6 | 17 | 15 | 63 | 7 | 44 | 11 |
| BA | 0.021 | 0.099 | 0.000 | 0.027 | 0.078 | 0.112 | 0.098 | 0.173 | 0.095 |

To visualize overall dietary similarity among species, we estimated a non-metric multidimensional scaling (NMDS) plot from a Bray-Curtis dissimilarity matrix of dietary proportions for each individual. To test for differences in diet among species, we used analysis of similarities (ANOSIM) with species designated as the grouping variable. To determine which dietary components significantly contributed to the stomach contents of each species, we performed an indicator species analysis (Defrêne & Legendre 1997; Cácaeres & Legendre 2009). This analysis has traditionally been used to identify one or more species characterizing various habitats or sites in ecological studies (Defrêne & Legendre 1997). More recently, it has been used in dietary studies to identify diet items significantly contributing to differences in stomach contents between groups (Hertz *et al.* 2017; Lee *et al.* 2018; Thalmann *et al.* 2020). These visualizations and analyses were performed in R using the *vegan* (Oksanen *et al.* 2019) and *indicspecies* (Cáceres & Legendre 2009) packages.

To determine whether individual dietary proportions varied by species, we used generalized linear models (GLMs). All GLMs were fitted using the *stats* package in R (R Core Team 2020). Dietary proportions were first transformed using the arcsine (also known as arcsine square root) transformation typical for proportional data. We fit a separate model for each dietary item after arcsine-transformation of the proportions. The independent variable was species with log-transformed standard length (SL) as a covariate. A normal distribution was used for all models. To test the significance of each model, we performed an ANOVA with Type III sum of squares using the *car* package (Fox & Weisberg 2019) in R. For significant models, we used Tukey’s HSD post hoc analysis for pairwise comparisons between species. Post-hoc analyses were conducted using the *stats* package in R.

Since volume-based dietary proportions are highly variable depending on prey condition (Buckland *et al.* 2017), we decided to additionally analyze our stomach content data using a frequency of occurrence approach. To determine whether presence/absence of dietary components varied by species, we fit GLMs using a binomial distribution. The proportional response variable was converted into binomial success (proportion > 0) and failure (proportion = 0) and then all models were fit as described above.

To determine whether stable isotope content varied by species, we fit GLM models for both δ^15^N and δ^13^C. We fit models including all 11 Barombi Mbo cichlids and models including only the three *Stomatepia* species. The response variable for each was δ^15^N and δ^13^C, respectively, and the independent variable was species. A normal distribution was used for all models. To test the significance of each model, we performed an ANOVA with Type III sum of squares. We used Tukey’s HSD post hoc analysis for pairwise comparisons between species.

## RESULTS

### Dietary composition and niche breadth

We found a majority of Barombi Mbo cichlids consumed detritus, plant tissue, and aquatic insects (Fig. 2). Detritus was the majority (>50%) dietary component in all species except for *S. mongo* (Table 1; Fig. 2). Notably, *Myaka myaka* was the only species with stomach contents consisting of 100% detritus (Table 1), potentially reflecting rapid digestion of its specialized *Chaoborus* larvae diet (Trewavas et al. 1972) into unidentifiable organic matter or capture in 2016 during the summer lekking season when reproductive males may invest all their time in courting females rather than foraging. *Konia eisentrauti* consumed the largest percentage of plant tissue (22.8%) across all species (Table 1; Fig 3). While *Pungu maclareni* and all three *Stomatepia* species consumed shrimp, *Stomatepia mongo* consumed the greatest proportion of shrimp (62.9%) among all species (Table 1; Fig. 1 & 3). This may reflect the rare hunting strategy of this species for nocturnal shrimp prey (also see Lloyd *et al.* 2021 for a nocturnal Malawi cichlid). *S. mongo* were only observed and captured by seine net after twilight hours beginning around 19:00 hours. Most species also had unidentified material in their stomach contents, although this percentage was typically under 15% on average (Table 1). However, *S. steinbachi* contained the highest percentage (38.3%) of unidentified material (Table 1), specifically egg-like structures that could not be identified.

**Figure 1:**
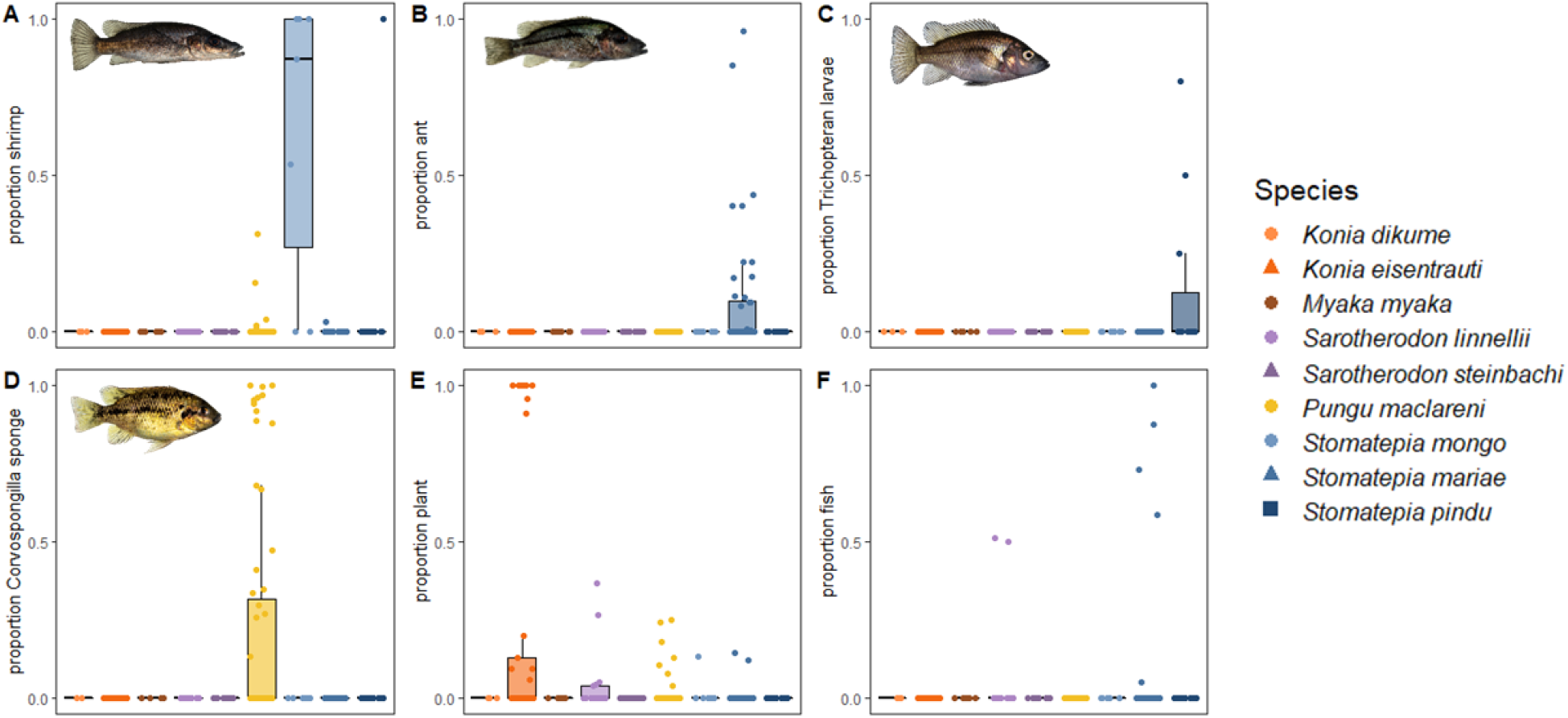
Box and whisker plots displaying the proportions of A) shrimp B) ants, C) Trichopteran larvae, D) *Corvospongilla* sponge, E) plant tissue, and F) fish found in each species’ stomachs. Total sample was 203 individuals collected in 2010 and 2016 from multiple sites around the lake.

**Figure 2:**
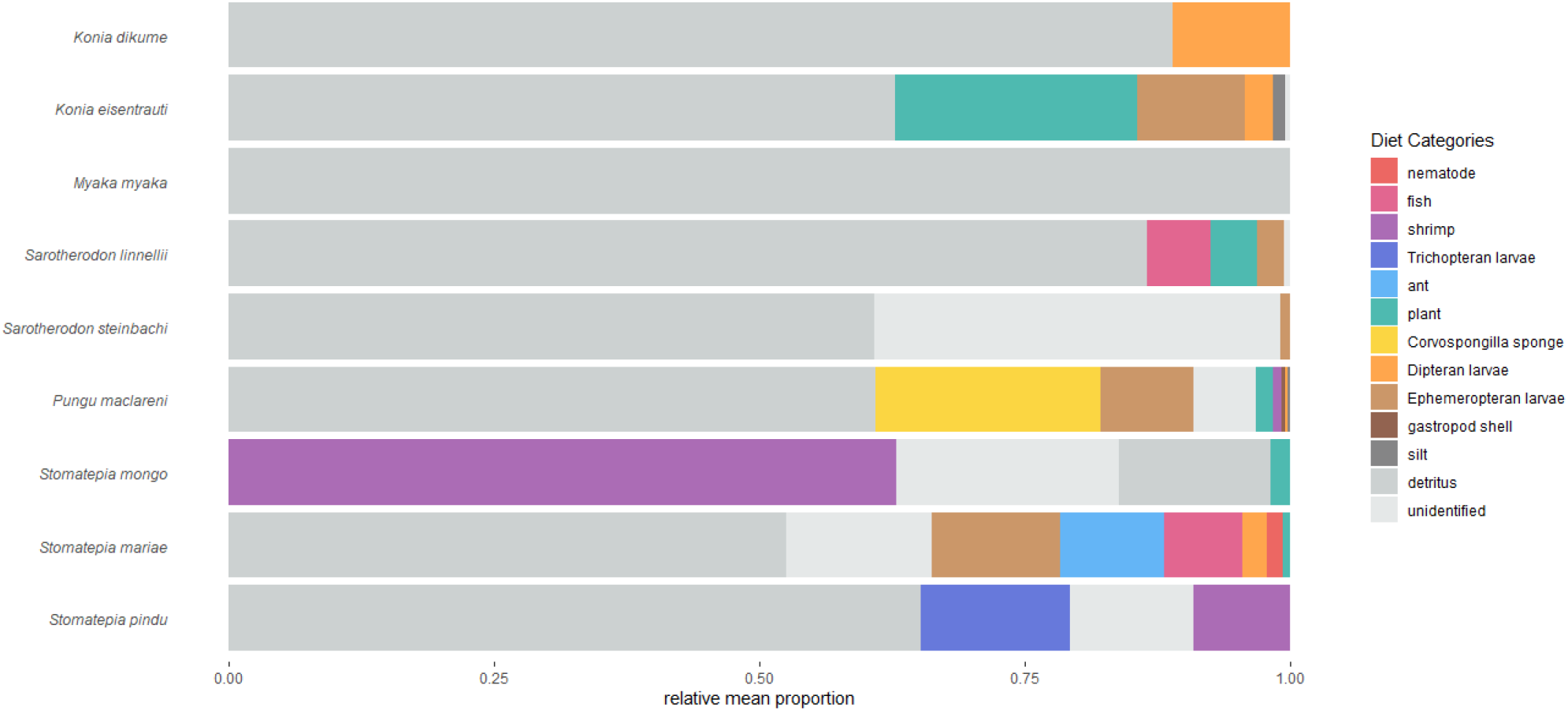
Dietary profiles of Barombi Mbo cichlids by prey item proportion. Each color represents a different dietary component. Bar length is based on the average proportion of each prey item.

**Figure 3:**
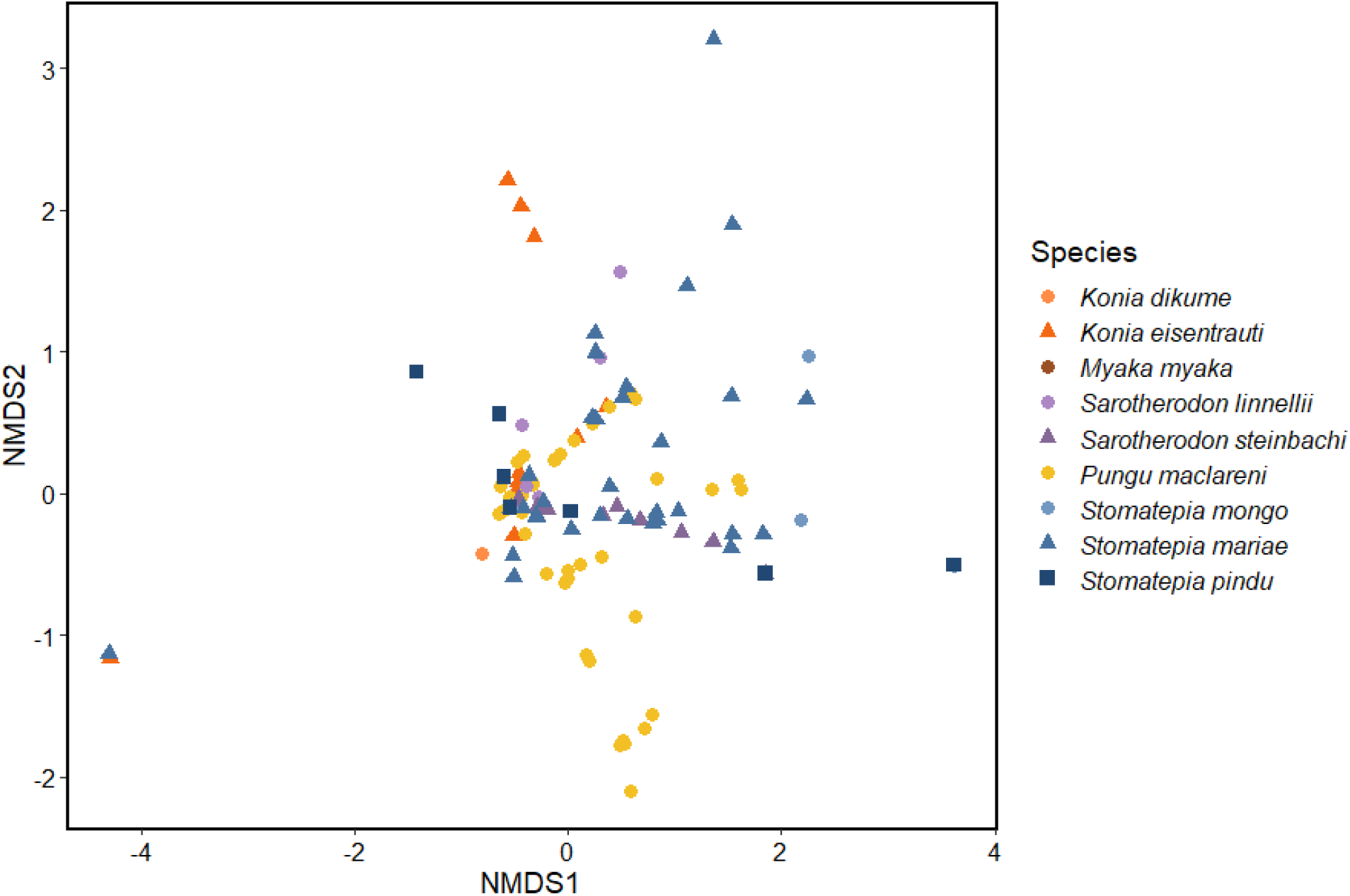
Non-metric multidimensional scaling (NMDS) ordination of dietary item proportions for nine out of the eleven Barombi Mbo cichlids. Ordination is based on Bray-Curtis similarity index (stress = 0.103).

Several dietary components were much rarer and found in only one or two cichlid species. *S. linnellii*, the largest species in the radiation, and *Stomatepia mariae* were the only two species to consume fish (Table 1; Fig. 1). *P. maclareni* was the only species to consume gastropod shells and *Corvospongilla* sponge spicules, with the latter component making up about 20% of this species’ diet on average (Table 1; Fig. 1). *S. mariae* was the only species to consume ants, comprising about 10% of this species’ diet on average (Table 1; Fig. 1). While insect larvae from the orders Ephemeroptera and Diptera were found in the stomach contents of several species, only *Stomatepia pindu* consumed Trichopteran larvae (Table 1; Fig. 2), comprising about 14% of this species’ diet on average (Table 1; Fig. 1).

*S. linnellii* had the widest niche breadth among all species, whereas *M. myaka* had the smallest (Table 1). Many species displayed considerable dietary niche overlap, with Pianka index values typically ranging from 0.8-1 (Table 2). Notably, *S. mongo* showed the lowest niche overlap with other species, with Pianka index values from 0.2-0.35 (Table 2), possibly reflecting the combination of both temporal and trophic niche divergence in this species.

**Table 2:** Pianka’s measure of niche overlap (Pianka 1973) among Barombi Mbo cichlid species. Values range from 0-1, with 0 being no niche overlap and 1 being complete niche overlap.

|  | *Konia dikume* | *Konia eisentrauti* | *Myaka myaka* | *Sarotherodon linnellii* | *Sarotherodon steinbachi* | *Pungu maclareni* | *Stomatepia mongo* | *Stomatepia mariae* |
| --- | --- | --- | --- | --- | --- | --- | --- | --- |
| *Konia eisentrauti* | 0.926 |  |  |  |  |  |  |  |
| *Myaka myaka* | 0.992 | 0.928 |  |  |  |  |  |  |
| *Sarotherodon linnellii* | 0.988 | 0.946 | 0.996 |  |  |  |  |  |
| *Sarotherodon steinbachi* | 0.840 | 0.791 | 0.846 | 0.847 |  |  |  |  |
| *Pungu maclareni* | 0.925 | 0.894 | 0.932 | 0.934 | 0.838 |  |  |  |
| *Stomatepia mongo* | 0.209 | 0.207 | 0.211 | 0.213 | 0.342 | 0.236 |  |  |
| *Stomatepia mariae* | 0.919 | 0.894 | 0.921 | 0.935 | 0.910 | 0.909 | 0.270 |  |
| *Stomatepia pindu* | 0.947 | 0.887 | 0.954 | 0.951 | 0.899 | 0.906 | 0.377 | 0.920 |

### Clustering, analysis of similarities, and indicator analyses of overall diet

The NMDS ordination (stress = 0.103) displayed little clustering of species by dietary components, with considerable overlap among species (Fig. 3). However, there was a statistically significant difference in overall diet among species (ANOSIM: R = 0.06275, *P* = 0.0238).

We identified several dietary items that significantly predicted species identity. Detritus was a significant indicator of *K. dikume, M. myaka*, and *S. linnellii* (indicspecies: Dufrêne-Legendre indicator value = 0.454, *P* = 0.0097). Plant tissue was a significant indicator of *K. eisentrauti* (Dufrêne-Legendre Indicator = 0.438, *P* = 0.0199); shrimp was a significant indicator of *S. mongo* (Dufrêne-Legendre Indicator = 0.744, *P* = 0.0003); *Corvospongilla* sponge was a significant indicator of *P. maclareni* (indicspecies: Dufrêne-Legendre indicator value = 0.49, *P* = 0.0153); ants were a significant indicator of *S. mariae* (indicspecies: Dufrêne-Legendre indicator value = 0.405, *P* = 0.0448); and Trichopteran larvae were a significant indicator of *S. pindu* (indicspecies: Dufrêne-Legendre indicator value = 0.457, *P* = 0.0271).

### Individual dietary components

We found individual diet proportions to vary by species for several items (Fig. 1). Detritus consumption was significantly different among species (ANOVA: χ2 = 36.242, df = 8, P = 1.585×10-5). In particular, *M. myaka* consumed about 2 times more detritus than *S. mariae* (Tukey HSD: *P* = 0.044), and *S. linnellii* consumed about 1.5 times more detritus than *S. mariae* (Tukey HSD: *P* = 0.021). Plant tissue consumption varied across species (Fig. 1E; ANOVA: χ^2^ = 49.347, df = 8, *P* = 5.455×10^−8^), with *K. eistentrauti* consuming at least 5 times more plant tissue than *S. linnellii*, *S. steinbachi*, *P. maclareni*, *S. mariae*, and *S. pindu* (Tukey HSD: *P* < 0.05). Shrimp consumption also varied among species (Fig. 1A; ANOVA: χ^2^ = 116.674, df = 8, *P* < 2×10^−16^), with *S. mongo* consuming at least 7 times more shrimp than all other species (Tukey HSD: *P* < 0.001). Consumption of unidentified diet items varied among species (ANOVA: χ^2^ = 55.175, df = 8, *P* = 4.082×10^−9^), with *S. steinbachi* consuming at least 2 times more than all other species (Tukey HSD: *P* < 0.05). *P. maclareni* was the only species to consume *Corvospongilla* sponge (Fig. 1D; ANOVA: χ^2^ = 55.461, df = 8, *P* = 3.591×10^−9^). *Corvospongilla* spicules made up 21.2% of *P. maclareni’s* diet (Table 1; Fig. 2). Similarly, *S. mariae* was the only species to consume ants (Fig. 1B; ANOVA: χ^2^ = 51.806, df = 8, *P* = 1.835×10^−8^). Ants made up 9.8% of *S. mariae’s* diet (Table 1; Fig. 2). *S. pindu* was the only species to consume Trichopteran larvae (Fig. 1C; ANOVA: χ^2^ = 58.100, df = 8, *P* = 1.098×10^−9^). Trichopteran larvae made up 14.1% of *S. pindu’s* diet (Table 1; Fig. 2).

Collapsing these proportional data to presence/absence data of individual dietary components (as described above) yielded similar results. Detritus (ANOVA: χ^2^ = 24.440, df = 8, *P* = 0.002), plant tissue (ANOVA: χ^2^ = 36.158, df = 8, *P* = 1.643*10^−5^), shrimp (ANOVA: χ^2^ = 19.1124, df = 8, *P* = 0.014), unidentified items (ANOVA: χ^2^ = 45.748, df = 8, *P* = 2.654*10^−7^); *Corvospongilla* sponge (ANOVA: χ^2^ = 55.628, df = 8, *P* = 3.333*10^−9^); ants (ANOVA: χ^2^ = 51.997, df = 8, *P* = 1.685*10^−8^); and Trichopteran larvae (ANOVA: χ^2^ = 17.6528, df = 8, *P* = 0.024), all varied significantly among species by presence/absence with similar specialists as described above.

### Carbon and nitrogen stable isotopes

We found δ^13^C values to be significantly different among species when comparing all 11 Barombi Mbo cichlids (ANOVA: χ^2^ = 123.36, df = 11, *P* = 2.2×10^−16^). *S. lohbergeri* had the highest δ^13^C value, indicative of predominantly littoral foraging, significantly more than all other species except *P. maclareni*, *S. caroli*, and *S. steinbachi* (Tukey HSD: *P* < 0.01). Contrastingly, *M. myaka* had the lowest δ^13^C value, consistent with its open-water pelagic habitat, significantly lower than all other species except *K. dikume*, *K. eisentrauti*, *S. linnellii*, and *S. pindu* (Tukey HSD: *P* < 0.05). *S. mongo* exhibited significantly higher δ^13^C than *S. pindu* when comparing all Barombi Mbo cichlids (Tukey HSD: *P* = 0.041) and only *Stomatepia* species (Tukey HSD: *P* = 0.005).

δ^15^N values were also significantly different among species when comparing all Barombi Mbo species (ANOVA: χ^2^ = 67.967, df = 11, *P* = 2.969×10^−10^). *K. dikume* had the highest δ^15^N value of any species, significantly more than *M. myaka*, *S. caroli*, *S. lohbergeri*, and *S. pindu* (Tukey HSD: *P* < 0.05). *S. lohbergeri* had the lowest δ^15^N value, significantly lower than all other species except *M. myaka*, *S. caroli*, and *S. steinbachi* (Tukey HSD: *P* < 0.05). There were no significant differences in δ^15^N values among *Stomatepia* species. Despite these significant differences in both δ^13^C and δ^15^N values between species, there was minimal clustering by species when visualizing stable isotope values (Fig. 4).

**Figure 4:**
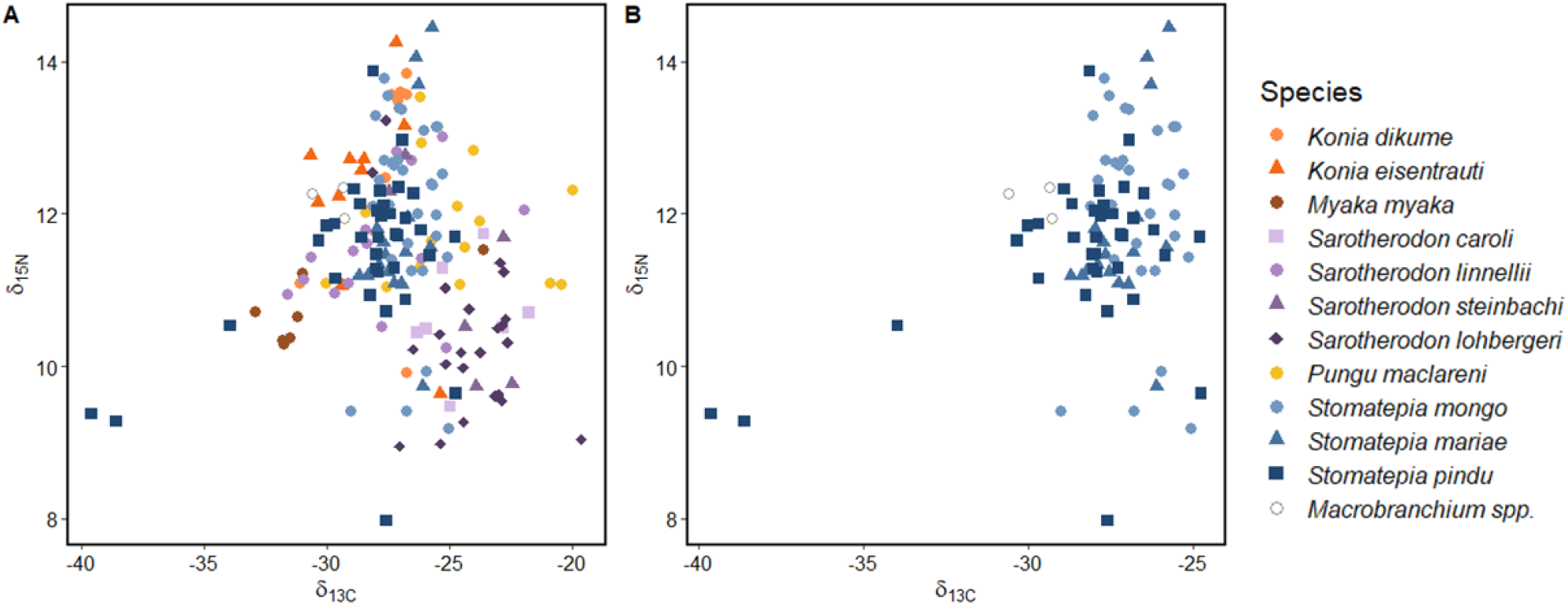
Scatterplots of δ^13^C and δ^15^N isotopic values for A) all eleven Barombi Mbo cichlids and B) only *Stomatepia* species. δ^13^C offers insight into ultimate carbon source (littoral vs pelagic) while δ^15^N values describe relative trophic position.

## DISCUSSION

We found minimal evidence of overall trophic niche partitioning among Barombi Mbo cichlids. However, we found several dietary specializations on unique resources among all freshwater fishes. In particular, our data suggests that Lake Barombi Mbo harbors a sponge specialist, *P. maclareni*, and an ant specialist, *S. mariae*. We also document a nocturnal specialist on shrimp (*S. mongo*) and specialists on Trichopteran (caddisfly) larvae (*S. pindu*), and plants (*K. eisentrauti*). Herbivory is common among African cichlids (Ribbink & Lewis 1982; Genner & Turner 2005), and *K. eisentrauti* was previously qualitatively described as a plant specialist (Trewavas *et al.* 1972). However, the remaining resource specializations are particularly rare among African cichlids. Nocturnality has only been documented once out of thousands of Malawi cichlids (Lloyd *et al.* 2021) and spongivory is only qualitatively reported from *Coptodon spongotroktis* in Lake Bermin, Cameroon (Stiassny *et al.* 1992). In general, our findings align with the major trophic strategies and specialists outlined by Trewavas *et al*. (1972), providing quantitative data on the differences in dietary component proportions between species.

### Minimal trophic niche partitioning in Barombi Mbo cichlids

Our measurements of niche overlap suggest there is not strong evidence for dietary niche partitioning in Barombi Mbo cichlids, with Pianka index values of 84% similarity and higher for most species. Furthermore, while we did find significant differences in δ^13^C and δ^15^N values between species, there was little evidence of clustering by species. This is not uncommon, as coexistence among ecologically similar species can occur even without fine-scale niche partitioning, particularly within speciose African cichlid communities in the great lakes (Liem 1980; Ribbink & Lewis 1981; Martin and Genner 2009).

Alternatively, dietary niche partitioning may have been obscured by variability in prey condition. Stomach content analyses are highly dependent on prey condition (Baker *et al.* 2014; Buckland *et al.* 2017). Soft-bodied organisms are likely to digest more quickly than those with chitinous exoskeletons or other similarly tough external features, potentially leading to an overrepresentation of hard-bodied, less digestible organisms (Randall 1967). Furthermore, there may be finer-scale niche partitioning at lower taxonomic prey levels (i.e. genus, species) than what can be detected by microscopic visual analysis of stomach contents. DNA metabarcoding approaches may aid in identifying these finer-scale patterns (Berry *et al.* 2015; Harms-Tuohy *et al.* 2016; Jakubavičiūtė *et al.* 2017) and can also account for highly digested prey (Carreon-Martinez *et al.* 2011), though such techniques come with their own suite of challenges and may overestimate the relative importance of certain dietary items (Sakaguchi *et al.* 2017).

Another explanation for the minimal trophic niche partitioning we observed could lie in undetected seasonal differences. Our data combined specimens collected during both the wet (July-September) and dry (December-January) seasons in Cameroon across multiple years (2009-10, 2016). Prey availability often differs greatly between seasons, especially in tropical systems (Winemiller 1990; Correa & Winemiller 2014). However, dietary profiles of the trophic specialists in Barombi Mbo were generally consistent between seasons and years collected. We conclude that these cichlids may be another example of Liem’s paradox, in which trophic specialists act as opportunistic feeders until resources become scarce enough that they use their specialized trophic morphology to feed on unusual resources. All specialists appear to be predominantly feeding on common, shared resources in Barombi Mbo (microinvertebrates, microalgae, detritus) while also supplementing their diet with unique resources during both the wet and dry seasons.

### Spongivory in *P. maclareni*

One of our most exciting findings is evidence of sponge specialization in *P. maclareni*. A significant proportion (20%) of this species’ diet is freshwater sponges (*Corvospongilla spp.*) and it is the only Barombi Mbo species to consume this diet item. Dietary specialization on sponges is extremely rare among fishes, with only about 0.04% of all fishes (FishBase) consuming sponges (2 out of a total of 13 entries are Cameroon crater lake cichlids). The most notable examples of this feeding strategy are marine spongivores that reside on Caribbean coral reefs (Randall & Hartman 1968; Wulff 1994). Eleven Caribbean reef fishes have been identified as sponge specialists, including angelfish in the genera *Holacanthus* and *Pomacanthus*, trunkfish in genus *Acanthostracion*, and filefish in genus *Cantherhines* (Randall & Hartman 1968). Sponge-eating is even more rare within freshwater systems, with the only two examples found in Cameroon crater lake cichlids of Lakes Barombi Mbo (Trewavas *et al.* 1972) and Bermin (Stiassny *et al.* 1992; Schliewen 2005). Compared to the proportions of sponge found in the stomachs of Carribean reef fish—*Holocanthus spp.* (>96%), *Pomacanthus spp.* (70-75%), *Cantherhines macrocerus* (86.5%) (Randall & Hartman 1968)—the proportion of sponge in *P. maclareni’s* stomach is small (20%), but still notable considering it is the only fish in this system to consume freshwater sponges. Furthermore, this proportion is still comparable to some of the other Carribean spongivores, including *Acanthostracion spp.* (11-30%) and *Cantherhines pullus* (30.9%) (Randall & Hartman 1968).

Sponges are a rare diet item among fish and other vertebrates because they are incredibly hard to consume. Most species in the Phylum Porifera have tough exteriors and skeletons made of spongin, calcium carbonate, and silica—all rigid materials. Spongivorous vertebrates, including the hawksbill turtle (*Eretmochelys imbricata*) (Meylan 1988; Witzel 1983; Eckert *et al* 1999) and several Carribean reef fishes have developed morphological adaptations in the feeding apparatus to aid in biting sponge spicules (Hourigan *et al.* 1989). In particular, *E. imbricata* possesses a narrow, beak-shaped mouth that allows for foraging on sponges in coral reefs (Witzel 1983; Eckert *et al.* 1999) and by scraping against the reef’s surface (Blumenthal *et al.* 2009). Several species of Carribean angelfish (*Holocanthus tricolor*, *Pomacanthus paru*, *Pomancanthus arcuatus*) also possess a beak-like mouth and multiple rows of tricuspid teeth used to shear sponge off its substrate (Hourigan *et al.* 1989). *P. maclareni* also appears to have adaptations that may aid in sponge-eating, including short robust oral jaws, large epaxial musculature (particularly when compared to other Barombi Mbo cichlids), and fleshy lips with protruding tricuspid teeth.

Spongivory in fishes may have evolved through modification of an algae-eating trophic strategy. Algivores often possess morphological and locomotory adaptations to aid in the biting, shearing, and scraping of algae attached to rocks and other substrates (Hulsey *et al.* 2019; Perevolotsky *et al.* 2020). This specialized feeding apparatus may have been co-opted and modified for a sponge spicule diet, as the functional and locomotory processes of shearing algae and sponge are likely similar. In fact, Caribbean spongivores use their peripheral teeth in a similar manner to tear both algae and sponge (Hourigan *et al.* 1989).

Another hurdle to spongivory lies in the difficulty of sponge digestion. Poriferans are typically made of materials (spongin, collagen, calcium carbonate, silica) that are difficult to digest and not nutritionally valuable to vertebrates. Furthermore, many sponges produce noxious secondary metabolites, including alkaloids, terpenoids, brominated compounds, and various acids (Faulkner 1984). Several marine sponges producing these compounds have been proven toxic to fish in lab experiments (Green 1977). To aid in these challenges, spongivorous fish may have morphological and physiological adaptations that allow for sponge digestion. Specifically, adaptations in the gut microbiome may be an essential component to sponge digestion. The gut microbiomes of other vertebrates have allowed for consumption and dietary specialization on rare food items, including vampire finches in the Galapagos (Michel *et al.* 2018; Song *et al.* 2019) and scale-eating pupfish (Heras & Martin 2021). Though the gut microbiome of wild-caught *P. maclareni* has been previously sequenced in a large comparative study (Baldo *et al.* 2017; Baldo *et al.* 2019), its functional relevance to spongivory was not assessed. Future research on *P. maclareni* and other spongivorous fishes should investigate the core gut microbiome of these species and any potential benefits the microbial community confers for sponge-eating.

### Ant consumption by *S. mariae*

Another notable finding of this study was evidence of ant specialization in *S. mariae*. We found that about 10% of this species’ diet is ants, and it is the only Barombi Mbo cichlid to consume this item. While *S. mariae* has previously been noted to feed on adult terrestrial insects (Trewavas et al. 1972), this is the first study documenting terrestrial ants as a major component of this species’ diet. There are several examples of freshwater fish in tropical systems in which ants have been observed as the majority dietary component, including flagtail *Kuhlia marginate* from Moorea, French Polynesia (Resh *et al.* 1999), and queen danio *Devario regina* from Malaysia (Zakeyuddin *et al.* 2017). Most relevant to this study are several species of Ecuadorian cichlids collected in the Upper Amazon River Basin (Saul 1975). All seven species collected (*Aequidens spp.*, *Crenicichla spp.*, *Petenia myseri*) consumed ants, with this food item being the most abundant dietary component in *C. lucius* and *C. macrophthalma* (Saul 1975). Ants were listed as an “occasional” food source for cichlids in this study (Saul 1975), a qualitative description that matches with our quantitative finding of ants making up 10% of *S. mariae’s* diet.

Terrestrial insects are not uncommon components of fish diets, as they enter lakes and rivers through fallen vegetation. The amount of terrestrial insects introduced into aquatic environments likely depends on vegetative and riparian cover, with insect abundance likely increasing as canopy cover increases (Bojsen 2005; Zakeyuddin *et al.* 2017). Fish skimming the water’s surface for food will take up these insects while foraging. This is likely why terrestrial insects are an important food source of known surface foragers (Resh *et al.* 1999; Nakano *et al.* 1999). While there have not been many studies on the subject, Sullivan *et al.* (2014) reported that the nutritional quality of terrestrial and aquatic arthropods is similar, particularly with respect to structural chitin—a limiting factor in nutrition as this polymer is prevalent in all arthropods and not easily digested by fish. Various species of terrestrial Cameroonian ants have been found to have high levels of protein (55-75% crude protein), and are rich in iron, zinc, potassium, phosphorus, and various other nutritionally-important minerals (Deblauwe & Janssens 2008). In fact, terrestrial insects, including ants, can subsidize the nutrient pool of small lakes, particularly those which have low primary productivity and are located in heavily forested areas (Mehner *et al.* 2005). Overabundance of terrestrial food sources in the diet of aquatic animals can indicate that the lake or stream is lacking in autochthonous nutrients (Saul 1975), as is the case for oligotrophic Barombi Mbo (Kling 1988).

It is interesting that *S. mariae* was the only Barombi Mbo cichlid to consume ants, as *S. mariae* and *S. pindu* are ecologically similar, hybridize in the lab, and represent the extreme tails of a unimodal distribution for all trophic traits measured (Martin 2012). The answer may lie in potential sensory adaptations that *S. mariae* uses to detect ants and its shoaling mid-water habitat, whereas *S. pindu* is a solitary benthic species that forages within the leaf-litter (Trewavas *et al.* 1972). Indeed, the genus *Stomatepia* was named for its highly enlarged canal neuromasts (stomae). Sensory traits of Barombi Mbo cichlids remain understudied, except for recent work showing differences in the visual sensory system and the pattern of photoreceptors among various species, with evidence of differences in opsin gene expression between shallow and deep-water species (Musilova *et al.* 2019). Future studies should investigate additional sensory systems in *S. mariae* and other Barombi Mbo specialists to better understand prey targeting of rare food sources.

## ACKNOWLEDGMENTS

This study was funded by a National Geographic Society Young Explorer’s Grant, a Lewis and Clark Field Research grant from the American Philosophical Society, the University of North Carolina at Chapel Hill, and the University of California, Berkeley to CHM. We gratefully acknowledge the Cameroonian government and the local chiefs of Barombi Mbo village and surrounding communities for permission to conduct this research. We also thank Jackson Waite-Himmelwright and Patrick Enyang for field assistance.

## Notes

### Competing Interest Statement

The authors have declared no competing interest.

